# Bayesian coalescent inference of in-host evolution using Next Generation Sequencing

**DOI:** 10.1101/407965

**Authors:** Gayle Leen, Marc Baguelin

**Affiliations:** Anthony Nolan Research Institute, London, United Kingdom; University College London Cancer Institute, London, United Kingdom; Public Health England, London, United Kingdom; London School of Hygiene and Tropical Medicine, London, United Kingdom

## Abstract

Within an infected individual, influenza virus exists as a heterogeneous population of variants. When representing the viral population as a consensus sequence, information about minority variants is lost. However, using next generation sequencing (NGS), it is possible to identify nucleotide substitutions which segregate at low frequencies in the viral population, and can give insight into the within-host processes that drive the virus’s evolution, and is a step towards understanding the dynamics of the disease. During the course of an infection, mutations may occur, and at each segregating site, the frequency of the derived allele in the population will fluctuate. We develop a method which can use information about the relative frequencies of mutations in NGS data from a viral population sampled at multiple time points, to infer past population dynamics with a Bayesian skyline model. By using coalescent theory, we analytically derive the joint allele frequency spectrum for a population across multiple time points, and relate this to the coalescent intervals generated from the skyline model. We demonstrate the model on data taken from populations of equine influenza virus sampled during an infection, and show that it is possible to infer a posterior distribution of effective viral population size through time. We also show how the model can be used to infer the probability that a mutation occurred within-host, as opposed to being an ancestral mutation which occurred prior to infection.

**Author Summary:** When a host is infected by a virus, many particles of the infecting agent enter the body of the host. This viral population is composed of many closely related viruses that continue diversifying by mutating while reproducing in the host. New sequencing technologies allow the quantifying of the proportion of the different variants present in the host at a particular time. Unfortunately, the data resulting from such sequencing techniques are difficult to interpret as they consist of many unlinked copies of relatively small fragments of genetic code distributed along the genome of the virus.

We designed a method combining models of virus genealogies and frequency of mutations appearing in the data to reconstruct the variation of the viral population inside the host. It also allows us to time the apparition of particular variants. This could be useful to detect if a particular mutation (e.g. providing drug resistance) has appeared in host or was circulating before. We applied our method to data of within-host evolution of equine influenza.

## Introduction

Understanding the evolution and kinetics of the within-host viral population during an influenza infection is an essential step to understanding the evolution and dynamics of viral populations on a larger scale. This is particularly important as viral evolution will have an impact on the dynamics of the disease, and potentially its virulence or resistance to some of the existing treatments. Influenza viruses exist as heterogeneous populations of evolutionarily related haplotypes in their infected hosts, due to short viral generation times and high mutation rates. In a within-host population, the genetic diversity is shaped at both the between-host level, due to host to host transmission dynamics and transmission bottlenecks, and the within-host level, through the host immune response, and the within-host demographic history.

Maintaining a diverse population of variants may be beneficial in adapting to such a dynamic environment, because a sudden change in environment which, in quasispecies theory terms, shifts the fitness landscape, may favour previously less fit, minority variants in maintaining the population’s survival [1]. However, it has been found that for influenza A virus, even though there is rapid variation over time (on a scale of years), there is low genetic diversity within a single host, and low genetic diversity across the global population at a single time [2]. Characterising the dynamics of the minority variants of the within-host viral population during an infection may be a step towards explaining this pattern.

Interactions between virus and host have previously been studied with viral kinetic models, where the characteristics of a viral growth and decay curve is governed by a set of biological parameters, and are fitted to viral load estimates sampled over time [3, 4]. However, the relationship between viral load and genetic diversity is unclear. Phylodynamics methods based on concepts from the population genetics field can be used to make inferences about the changes in population size from the patterns of mutations among sequences sampled from within-host over time [5–7]. In general, these methods summarise samples of the within-host viral population as consensus sequences, and use these to infer changes in effective population size, such as the HIV intrahost study in [8].

Characterising the sequence variation of a viral population has been enabled by recent advances in sequencing technologies, allowing researchers to look beyond the ‘consensus view’ [9]. Previously, information about the genetic diversity could only be accessed through isolating and cloning individual viruses and then applying Sanger sequencing [10, 11], which identified the predominant variants in the population. Next generation sequencing (NGS) technologies enable quick and low-cost sequencing of viral populations at high coverage, giving insight into the low frequency variants present in the sample. Though identification of the low frequency variants can be problematic, due to sequencing error rates, which may be of the same order as the variant frequency [12], and many existing variant calling algorithms focus on sites segregating at discrete frequencies, whereas in a viral population, the number of genomes, and hence the possible variant frequencies, are unknown, robust approaches to calling single nucleotide variants (SNVs) from high coverage datasets have been developed recently [12, 13].

Sequencing a viral population using NGS restricts genetic diversity estimation to SNVs or local fragments constrained by the read length. Though methods to assemble individual viral haplotypes have been proposed [14, 15], they can be computationally intensive. Consequently studying intrahost viral evolution at the level of whole genomes from NGS data is not straightforward, since existing phylodynamics methods e.g. [16–18] rely on having a known number of viral sequences. In this paper, we do not reconstruct the viral haplotypes; instead we analyse the viral population as a set of unlinked polymorphic sites (which we assume to be biallelic).

At each segregating site, we can calculate the frequency at which the derived allele appears, as the within-host viral population is sampled at various time points during an infection. This type of multidimensional SNP frequency data has been shown to be informative about the joint demography history of multiple populations [19, 20], and we follow this approach in this paper. We use the joint allele frequency spectrum (JAFS) as a measure of genetic diversity in the viral population shared across multiple time points, and assume that the population at each time point contains *N* genomes.

## Results

### New methodology for in-host evolution

We developed a Bayesian method for estimating the trajectory of effective viral population size over time, from next generation sequencing of the within-host viral population sampled at *P* time points. The genetic diversity in each sample is represented by a population of arbitrary size *N*, and the number of carriers of the derived allele *k*, (*k ≤ N*), is evaluated at a set of unlinked biallelic sites. For each site, the data is given as a segregating site pattern across the *P* time points.

We derived the probability of each pattern by using coalescent theory, following results derived for the joint allele frequency spectrum for multiple populations in [20]. The dependencies between the viral populations across the time points is modelled through a coalescent tree which represents the ancestral relationships between the *N×P* tips. We obtained a likelihood function for the data *D*, given the expected coalescent intervals 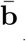 of the underlying genealogy 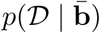.

We used the Bayesian skyline plot framework [16] to place a prior on 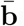. This quantifies the relationship between the length of the coalescent intervals of the genealogy of the viral population members, and the demographic history. As opposed to depending on a pre-defined parametric model, such as constant, or exponential growth, the skyline plot framework allows a flexible prior on possible histories, using a piecewise constant demographic model of effective population size through time. The model was parameterised by an ordered subset of group sizes A= {*a*_1_, *…, a*_*m*_}, which defines the number of coalescent events in each grouped interval, and *m ≤* (*N × P -* 1). Each of the *m* groups was assumed to have a constant effective population size Θ = {*θ*_1_, *…, θ*_*m*_}.

An MCMC scheme was used to sample from the posterior distribution 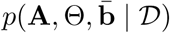. Given a set of posterior samples 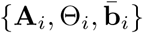, for each sample, we could evaluate the marginal posterior distribution of *θ*(*t*) at a set of times of interest *t*_*j*_, such that we have a set of marginal distributions *θ*(*t*_*j*_), that represent the estimated posterior effective population size through time, with its uncertainty.

### Simulated data

To investigate the behaviour of our intrahost model, we analysed simulated data sets from a basic model of viral dynamics (see Appendix). We fixed some of the parameters (*δ, c, k, T*_0_) to average values obtained from equine influenza infections in [4], as in Table 1, and set *V*_0_ = 10. We generated sets of data for 3 scenarios: {*β* = 1.4*×*10^−4^, *p* = 1.4*×*10^−5^}, {*β* = 1.4*×*10^−4^, *p* = 4*×*10^−6^},{*β* = 1*×*10^−4^, *p* = 2.8*×*10^−6^}, which have a viral peak at around 2, 3, and 4 days respectively. Each data set was generating using the following procedure: (1) The viral trajectory was generated given the parameters. (2) A coalescent tree was generated for *N* sequences taken at time 2, 4, and 6 days (3 * *N* sequences in total), by generating the coalescent intervals as described in the previous sections, and then generating a random topology. (3) *M* segregating sites were generated. A mutation event was generated by picking a branch of the tree with probability proportional to the branch length, and all descendants of the branch shared the mutation. Counts of the number of sequences at each sampling point which shared the mutation were stored e.g. {*k*_1_, *k*_2_, *k*_3_}

**Table 1.** Parameters for model of intrahost viral dynamics

| Parameter | Definition | Average value from [4] |
| --- | --- | --- |
| $\beta$ | Infection rate constant | $1.4 \times 10^{-4} [\text{TCID}_{50}/\text{ml}]^{-1} d^{-1}$ |
| $V_0$ | Initial virus titer | $9.3 \times 10^{-2}$ |
| $p$ | Average rate of increase of viral titer/infected cell | $1.4 \times 10^{-5} [\text{TCID}_{50}/\text{ml}] d^{-1}$ |
| $\delta$ | $1/\delta$ is the infected cell lifespan | $2d^{-1}$ |
| $c$ | Viral clearance rate | $5.2d^{-1}$ |
| $k$ | $1/k$ is the transition time between $I_1$ and $I_2$ | $2d^{-1}$ |
| $T_0$ | Initial number of uninfected target cells | fixed at $3.5 \times 10^{11}$ |

We ran the MCMC inference for each data set using a burn-in of 100, 000 samples, and for a chain of 2, 000, 000 samples, with thinning of 100, with 200 segregating sites, and *N* = 50. We used *m* = 15 population size intervals. Figure 1 shows the posterior viral trajectories from the model. The model manages to capture the growth in population size after infection at *t* = 0, and performs better when the rate of increase is slower i.e. when the population peaks at 4 days (right plot). We also find that the model does not capture the decrease in viral population after the peak. This is due to the large coalescent intervals just before the last sample at 6 days (long terminal branches in the genealogy), which are uninformative about change in population size.

**Figure 1.**
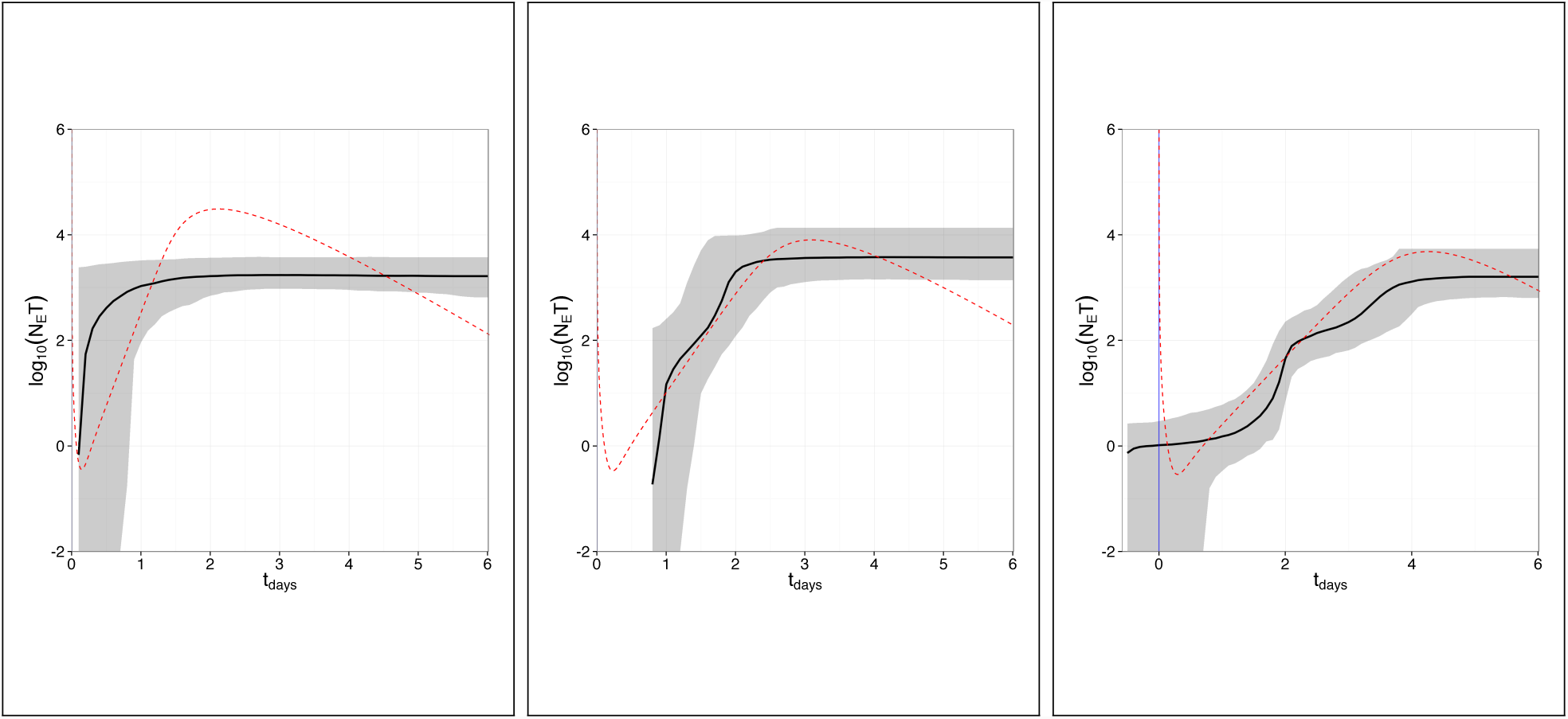
Performance of model on simulated data. Data sets of 50 samples at times 2, 4, and 6 days (150 tips in total), with 200 segregating sites, were generated from coalescent intervals under a viral dynamical model to given effective viral population size trajectories peaking at 2 (left), 3 (middle), and 4 (right) days. The true product of the effective viral population size and generation time (*N*_*e*_*τ*) is shown as a dashed red line. The median (black) and 95% confidence intervals (grey) of *N*-*eτ* estimated from the posterior samples from our model are summarised in a Bayesian skyline plot, where we have used *m* = 15 intervals.

### Intrahost evolution of equine influenza

We tested the model on intrahost viral populations sampled at multiple time points from horses experimentally infected with influenza A(H3N8) virus. [data available on request]. A majority consensus sequence was constructed from the inoculum sample, and used as the reference sequence. Daily samples from the horses were paired-end sequenced using the Illumina platform, and aligned with the Tanoti reference assembler http://www.bioinformatics.cvr.ac.uk/Tanoti/index.php. Quality checks were performed with FastQC http://www.bioinformatics.babraham.ac.uk/projects/fastqc/, followed by trimming of adaptors and low quality scores with Trim Galore! http://www.bioinformatics.babraham.ac.uk/projects/trim_galore/, resulting in a BAM file for each viral population at each time point. Duplicates were removed using Picard v1.119. Variants were called using LoFreq [13]. For each horse, we obtain a set of segregating sites, and the frequency of the derived allele at each site, across all time points. After assuming that the viral population can be represented by *n* = 50 viral genomes at each time point, and using (1), we obtain a set of segregating site patterns for each horse. The data sets consist of horses 2761 (N) (2, 3, and 5 days after infection), 6652 (2, 3, 4, 5), 6292 (3, 4, 5, 6), 1420 (3, 5, 6, 7), 2220 (4, 5, 6, 7), and 6285 (3, 5, 7).

For the data set from each horse, we ran the MCMC inference (burn-in of 100, 000 samples, for a chain of 2, 000, 000 samples, with thinning of 100) to estimate the effective population size trajectories. For all experiments, we use *m* = 15 population size intervals, and *N* = 50. Figure 2 shows the results from running the model on the data for the 6 horses. Each subfigure contains 3 plots. The top plot shows the posterior scaled effective population size trajectories (*N*_*e*_*τ*), with the median and 95% confidence intervals. The middle plot shows the posterior density of *p*(*t*_*µ*_) (red), the time at which the mutation arose, for each segregating site pattern. The densities are superimposed over each other, such that the darkest areas correspond to times where mutations are more probable. The mean of the densities is shown in black. The bottom plot shows the calculated number of viral copies per *µl*, obtained through qPCR. For all plots, the vertical blue lines show the start of infection (solid) and the times at which the viral population was sampled (dashed). The time axis is truncated to the median of the sum of branch lengths (tree height).

**Figure 2.**
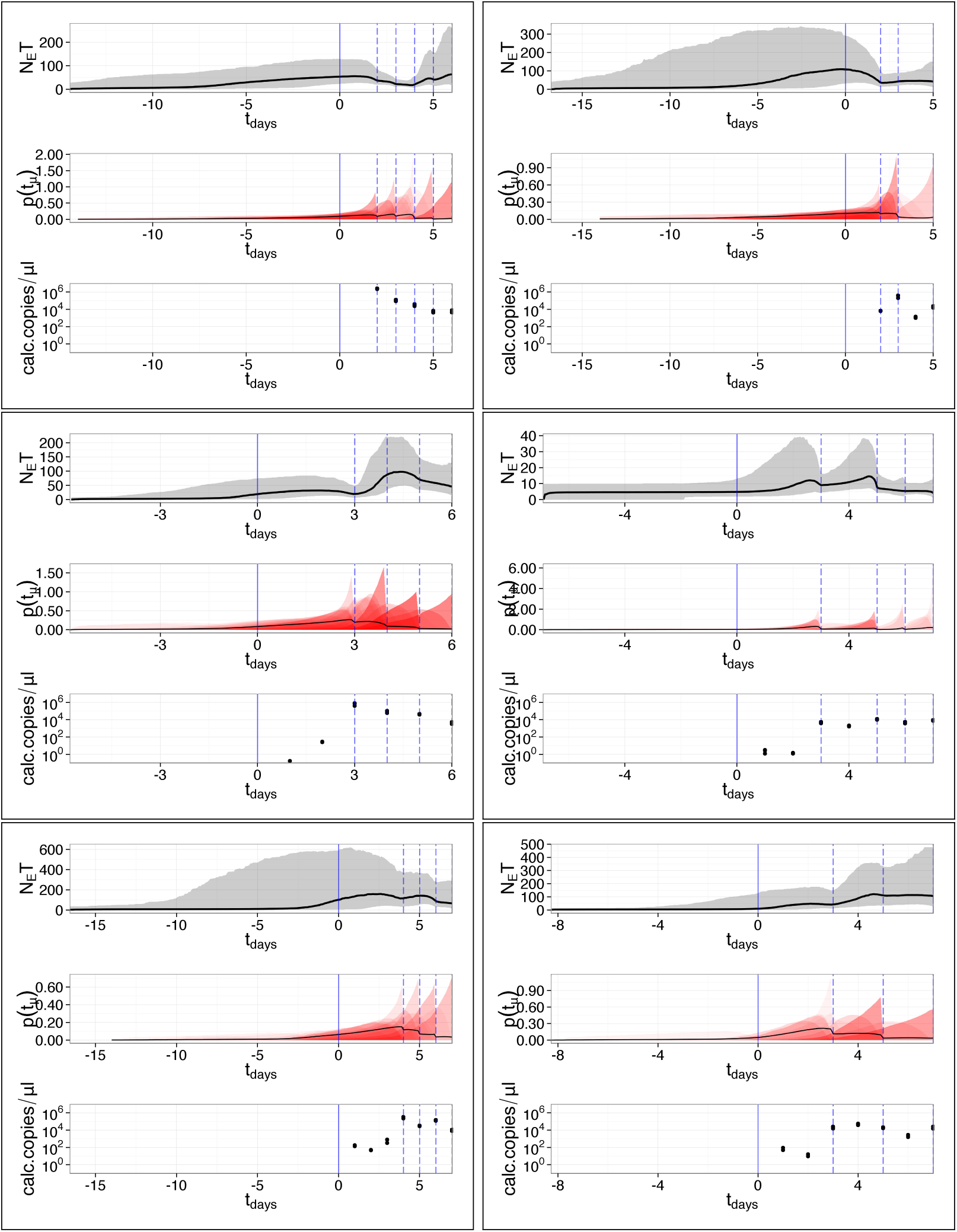
Effective population size trajectories. From left to right, top to bottom: horses 2761, 6652, 6292, 1420, 2220, 6285. Each subfigure contains 3 plots. The top plot shows the posterior scaled effective population size trajectories (*N*_*e*_*τ*), with the median and 95% confidence intervals. The middle plot shows the posterior density of *p*(*t*_*µ*_) (red), the time at which the mutation arose, for each segregating site pattern. The densities are superimposed over each other, such that the darkest areas correspond to times where mutations are more probable. The mean of the densities is shown in black. The bottom plot shows the calculated number of viral copies per *µl*, obtained through qPCR. For all plots, the vertical blue lines show the start of infection (solid) and the times at which the viral population was sampled (dashed). The time axis is truncated to the median of the sum of branch lengths (tree height).

For each mutation, we calculated the median probability that the mutation had risen within-host, *p*(*t*_*µ*_*≥*0). In Figure 3, we plot the frequency at which the mutation occurs in the viral population, against time (square points), for each horse. The trajectories (dotted line) are shaded according to probability *p*(*t*_*µ*_*≥*0), such that mutations estimated to have occurred within-host *p*(*t*_*µ*_*≥*0) are shaded purple, and those which were present before infection are shaded green.

**Figure 3.**
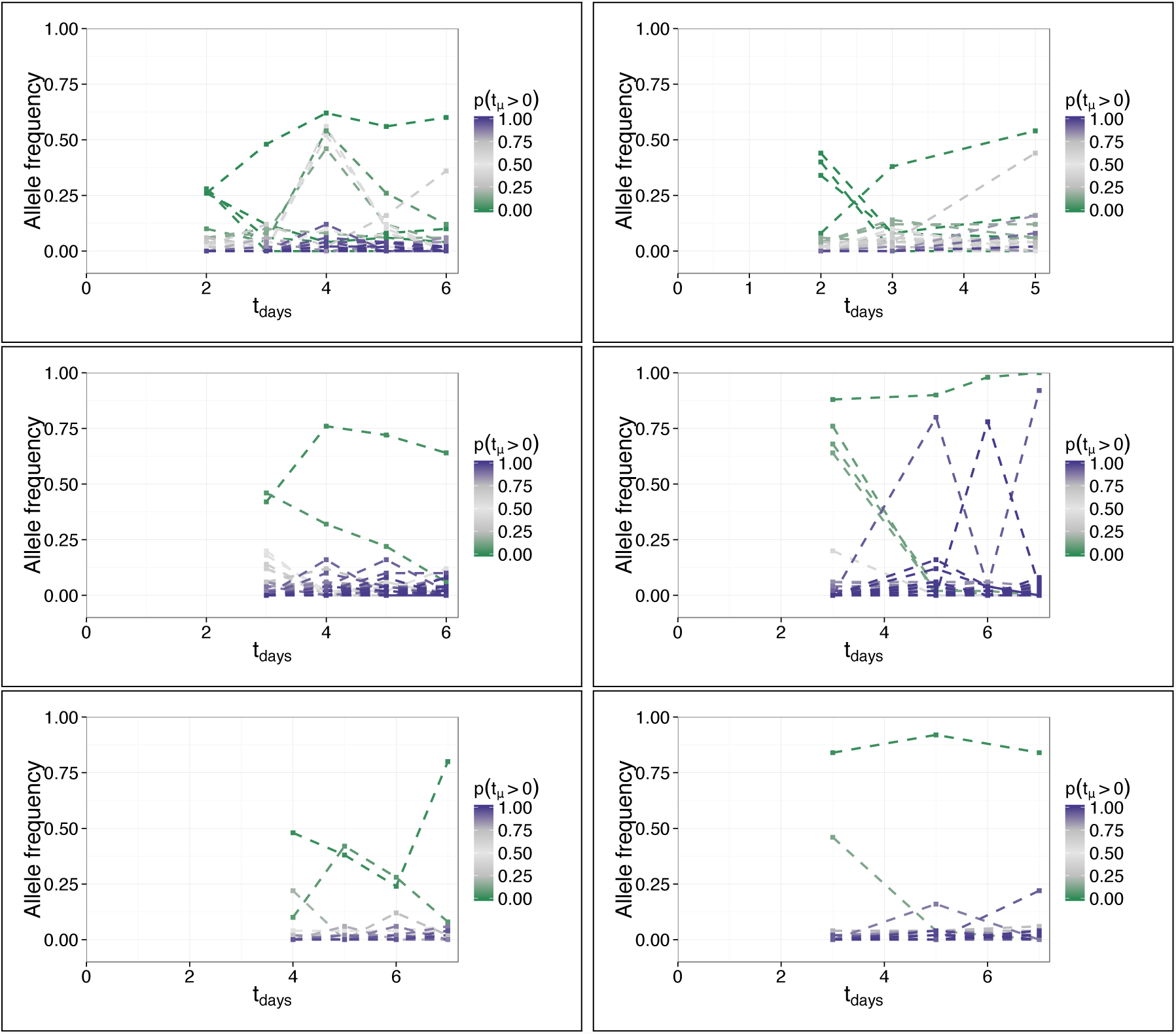
Allele frequency trajectories. From left to right, top to bottom: horses 2761, 6652, 6292, 1420, 2220, 6285. The frequency of each mutation in the viral population is plotted against time. The trajectories are shaded according to the probability that the mutation occurred within-host *p*(*t*_*µ*_*≥*0), such that the mutations most likely to have occurred within-host are purple, and those present before infection are shaded green.

## Discussion

Next generation sequencing of intrahost viral populations enables the identification of minority variants and their frequencies. Given NGS samples of a viral population over the course of an infection, we designed a novel methodology that uses the change in variant frequencies to infer in-host effective viral population size over time as a Bayesian skyline plot. The resulting skyline plots might be difficult to interpret as they relate to the effective viral population sizes rather than the actual number of viruses present in the host (viral load). This difficulty of interpretation also exists for the methodology as commonly applied at a (host) population level. Additionally, the resulting genealogy might present coalescent events which occurred before the time of infection. While the interpretation of what happens before the infection event is difficult, this reflects that diversity is present in a transmission event and passed from host to host. We are able to predict when particular mutations are likely to have appeared and thus inferring the probability of the mutation to have occurred in host. This could provide useful information while tracking the appearance of particular mutations such as the ones known to confer resistance to antiviral drugs.

While high frequency is usually an indicator of appearance outside of the host, it is not possible to predict from frequency only if a particular mutation has appeared inside or outside the host. Our methodology allows us to quantify the probability of both events. This would be particularly useful e.g. to study chains of transmission as low frequency minority variants might not appear in all samples or might be classified as error from the amplification process. Our study, by providing a tool to study in-host evolution of viral populations, paves the way toward the inference of chains of transmission where the additional mechanisms of loss of diversity through host-to-host bottlenecks would need to be integrated. Ultimately, this will lead to a better understanding of the drivers of evolution of viral populations during outbreaks [11].

## Materials and Methods

### Beyond the consensus

The ability to look beyond the consensus view of a heterogeneous viral population offers a way into characterising within-host sequence diversity during an infection, and to determine the population genetic processes which underlie the evolution of the population. Next generation sequencing (NGS) techniques can be used to resequence heterogeneous viral populations at high coverage, and reveal nucleotide substitutions present in a small fraction of the population. NGS, or high throughput sequencing estimates the sequence of an input sample by fragmenting it into short sections, or reads, which are then amplified and sequenced. The reads can then be aligned against a reference genome to determine if and where the input sample differs from the reference sequence.

When sequencing a viral population, the input sample consists of an unknown number of potentially different viral genomes. Because NGS requires the sample to be fragmented into short reads, it is challenging to determine from which genome a read originated, and consequently reconstruct the sequence of each individual genome. In this paper, we represent the genetic diversity of a viral population as a set of unlinked biallelic sites.

Suppose that we have sequenced a viral population over the course of an infection at time points *t*_1_, *…, t*_*P*_. For each of the *P* time points, we find the sites at which there are SNVs (relative to a reference sequence), and the frequency of the derived allele across the population. Let us denote the number of unique biallelic sites, which occur once or more across the *P* time points, as *S*. The observed allele frequency in the viral population over time for the *i*th segregating site is given by the vector q_*i*_ = {*q*_*i,*1_, *…, q*_*i,P*_}.

We assume that the sampled viral population at *t*_*p*_ is of size *n*_*p*_. The number of genomes that contain the *i*th segregating site over time is given by

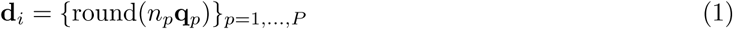

and the full dataset is given by *D* = {d_*i*_}_*i*=1,*…,S*_.

### Site frequency spectrum

The allele frequency spectrum (AFS) is a series of statistics that describe genetic polymorphism, and is defined as the sampling distribution of allele frequency at any random polymorphic locus in the genome. The AFS is informative for making inferences about the demographic history of a sample. While the majority of work has focused on using the AFS of a single population, advances have been made in using the joint allele frequency spectrum (JAFS) to infer more complex demographic events from multiple populations.

In this work, we derive the JAFS of multiple samples taken at different time points throughout an influenza infection, using coalescent theory. Given that we have *P* samples of an intrahost viral population, of sizes *n*_1_, *…, n*_*P*_ taken at times *t*_1_, *…, t*_*P*_, we denote the JAFS of the samples as m = {*m*_*k*1_,*…,k*_*P*_ (*n*_1_, *…, n*_*P*_), 0 *≤ k*_1_ *≤ n*_1_, 0 *≤ k*_2_ *≤ n*_2_, *…*, 0 *≤ k*_*P*_ *≤ n*_*P*_}, where *m*_*k*1_,*…,k*_*P*_ (*n*_1_, *…, n*_*P*_) denotes the probability that a SNP will have *k*_*i*_ copies of the derived allele in the *n*_*i*_ haplotypes of the *i*th population, *i* = 1, *…, P*.

We can derive the JAFS based on the genealogy underlying the *P* samples, with tips at *t*_1_, *…, t*_*P*_. Figure 4 shows an example genealogy underlying 3 samples of a population. Given the branching times of the genealogy, we know the number of ancestral lineages at any time point. If we look at time going forwards, and the first time of sampling *t*_1_, we note that *n*_1_ lineages are sampled, while the unsampled, remaining lineages are the ancestors, at *t*_1_, for the remaining tips of the tree. This continues along the tree, until the last sampling event at *t*_*P*_. If a mutation occurs along a branch of the genealogy, it is passed down via the branch’s descendants to the tips. (see Figure 5). The pattern of the segregating site across the tips of the genealogy is determined by the topology. It is possible to integrate over all the possible topologies to analytically find the probability of a segregating site pattern, following ideas in [20, 21]. We summarise the required distributions in the following section:

**Figure 4.**
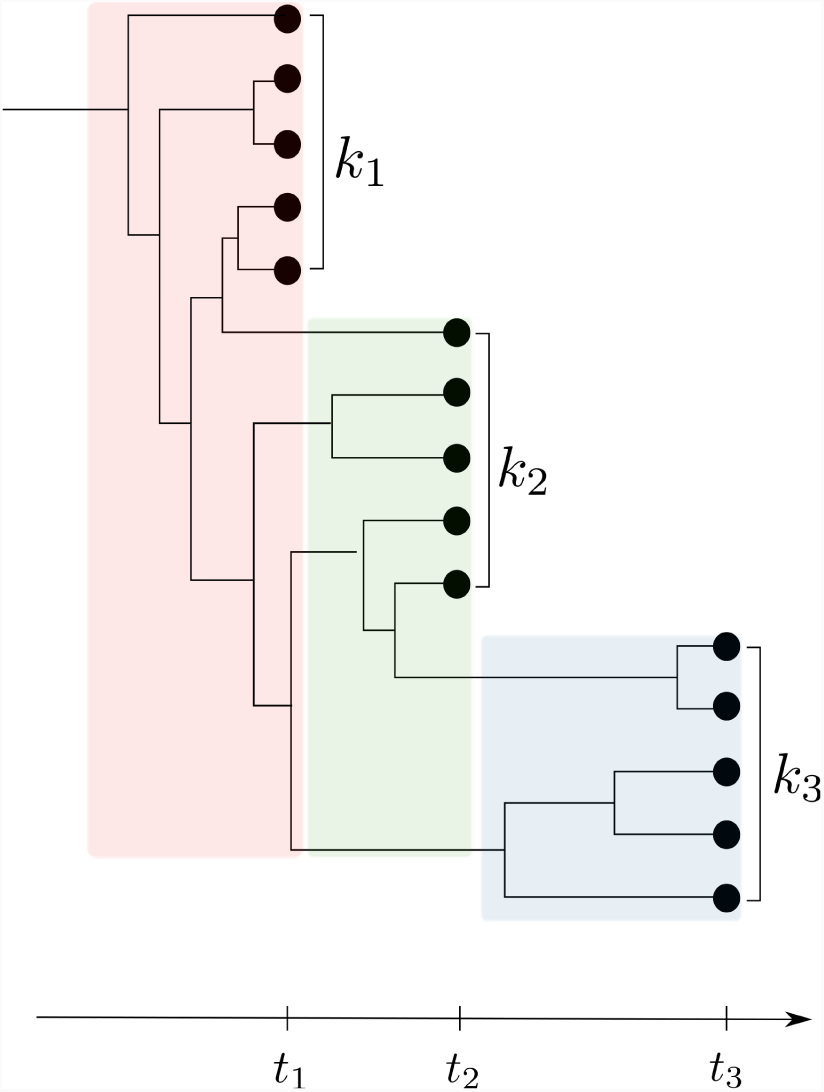
Possible genealogy underlying 3 samples. *k*_1_, *k*_2_, and *k*_3_ lineages are sampled from the population at times *t*_1_, *t*_2_, and *t*_3_. After sampling the population at *t*_1_, the unsampled lineages are the ancestors of the tips at *t*_2_ and *t*_3_, and similarly at *t*_2_.

**Figure 5.**
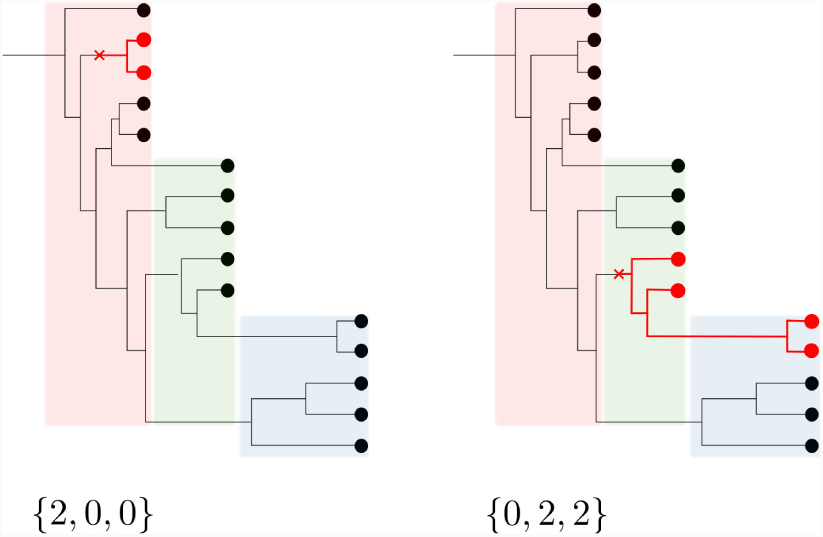
Possible genealogy underlying 3 samples with segregating site patterns from a mutation. When a mutation (shown in red) occurs in a lineage, its descendants inherit the derived allele. The number of copies of the derived allele in the sampled population at times *t*_1_, *t*_2_, and *t*_3_ are shown under each figure.

If a mutation occurs when there are *k* ancestors of a sample of *n*, then the probability that the mutation is found in *i* descendants in the *n* lineages is given by:

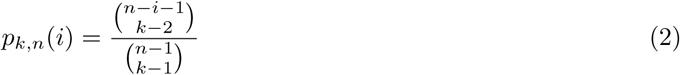

Suppose that there are *n* lineages, and *i* of these are carrying the same mutation. If the *n* lineages are split into two samples of size *m*_1_ and *m*_2_, (*m*_1_ + *m*_2_ = *n*), then the probability that there are *i*_1_ carriers in *m*_1_ and *i*_2_ in *m*_2_ (*i*_1_ + *i*_2_ = *i*), is given by the hypergeometric distribution:

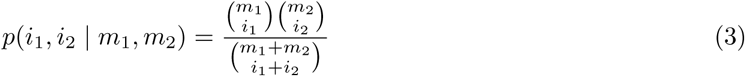

If there are *k* carriers of a mutation in *m* lineages, then the probability that, given that the *m* lineages grow to *n*, there will be *i* copies among the *n*, is given by the Polya-Eggenberger distribution:

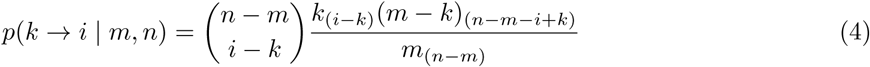

where *n*_(k)_ denotes the rising factorial 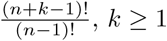.

#### Probability of a segregating site pattern

Given the *P* samples of the population (Figure 4), we calculate b, the expected lengths of the coalescent intervals of the underlying genealogy. We divide the timescale into stages, where the *i*th stage corresponds to the time between *t*_*i-*1_ and *t*_*i*_, the time points where the *i*th sample is taken from the population, *t*_0_ = 0. Each stage is further divided by the coalescent events that happen between its endpoints. We denote these intervals as 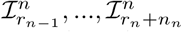 (in increasing chronological order), where 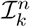 denotes the interval that contains *k* lineages, in the *n*th stage. The expected length of this interval is 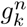. We denote the number of unsampled lineages at time *t*_*n*_ as *r*_*n*_. See Figure 6 for a schematic illustration of these terms.

**Figure 6.**
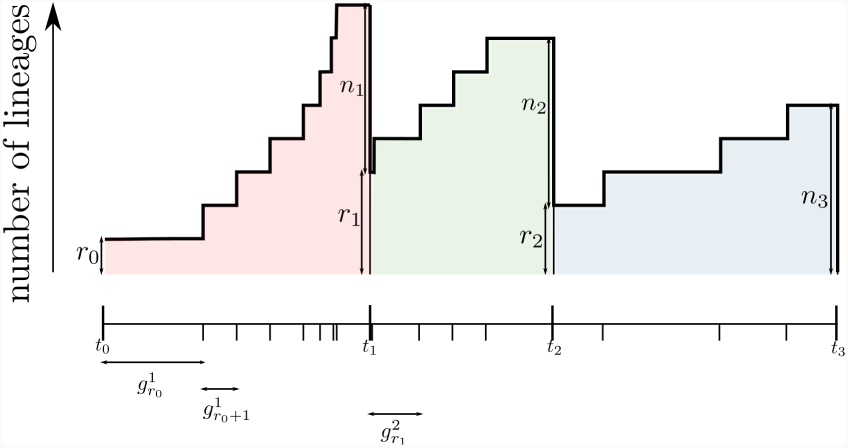
Change in the number of lineage for viral population sampled at multiple timepoints. At time *t*_*n*_, *n*_*n*_ lineages are sampled, and the remaining *r*_*n*_ lineages are the ancestors for the remaining tips in the tree. The expected length of the interval that contains *k* lineages in the *n*th stage (between *t*_*n-*1_ and *t*_*n*_) is denoted by 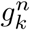.

At time *t*_*n*_, there are *n*_*n*_ sampled lineages and *r*_*n*_ unsampled lineages. Given that a mutation occurs between *t*_*n-*1_ and *t*_*n*_, the probability that there are *i*_*n*_ copies of the derived allele in the *n*_*n*_ sampled lineages and *j*_*n*_ copies in the *r*_*n*_ unsampled lineages is given by:

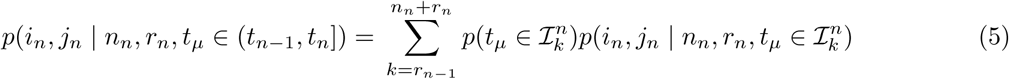

With

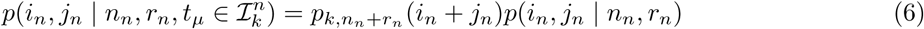

and where we have used (2) and (3), and 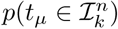 is the probability that the mutation occurs at stage *n*, when there are *k* lineages:

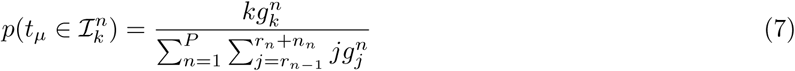

i.e. the branch lengths in 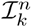 as a fraction of the total branch length of the tree.

The probability that the *j*_*n-*1_ derived alleles at time *t*_*n-*1_ have *i*_*n*_ descendants in the observed sample of size *n*_*n*_, and *j*_*n*_ of the *r*_*n*_ unsampled lineages at time *t*_*n*_ is given by:

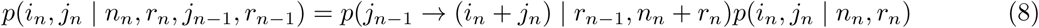

where we have used (3) and (4). We calculate the probability of a segregating site pattern {*i*_1_, *…, i*_*P*_} by summing over all possible configurations of {*j*_1_, *…, j*_*P*_}, the number of copies of the derived allele which are unsampled at {*t*_1_, *…, t*_*P*_}, and summing over all possible mutation events. The probability of {*i*_1_, *…, i*_*P*_} given that a mutation occurs in stage 1 is given by:

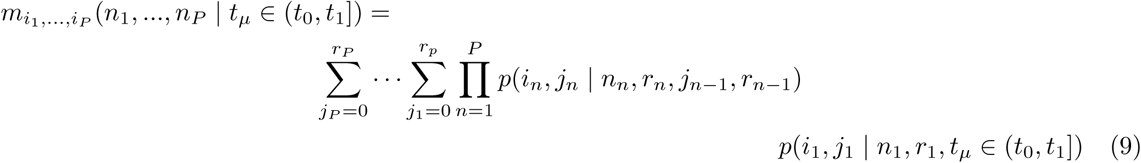

To include the probability of mutations in all the stages, we note that if we observe a site pattern where *i*_1_ *>* 0, the mutation will have occurred during stage 1, since it must have occurred in one of the ancestors of the *n*_1_ haplotypes that are sampled. More generally, we constrain the possible mutations to occur in stages before the first non-zero value in *i*. The probability of *i*_1_, *…, i*_*P*_ given that a mutation occurs in stage *p, p* = 2*…P*, is given by:

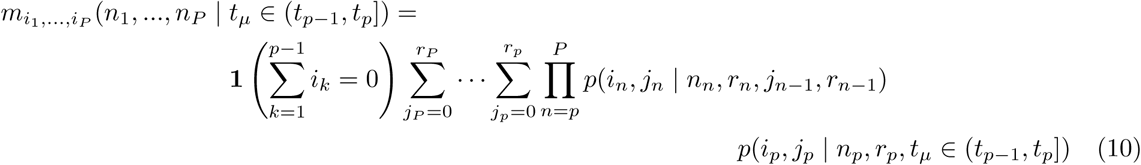

where 1() is the indicator function. The probability of the site pattern is given by the sum over all possible mutations

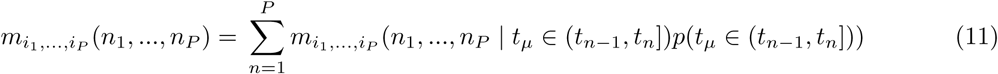

<u>a</u>nd we write the probability of a set of segregating site patterns *D*, given the average coalescent intervals 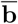of the underlying ancestral tree as

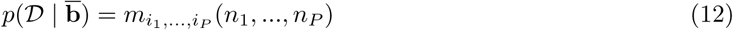

#### Age of a mutation

In the previous section, we derived the probability of a segregating site pattern, given the coalescent intervals of the underlying ancestral tree of the sample. We can estimate the age of the mutation which generated the site pattern d_*i*_ = {*i*_1_, *…, i*_*P*_}, by finding the posterior distribution of *t*_*µ*_. The probability that the mutation occurred during the *p*th stage (between *t*_*p-*1_ and *t*_*p*_), when there were *k* lineages is given by:

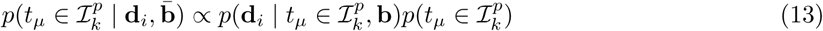

Where

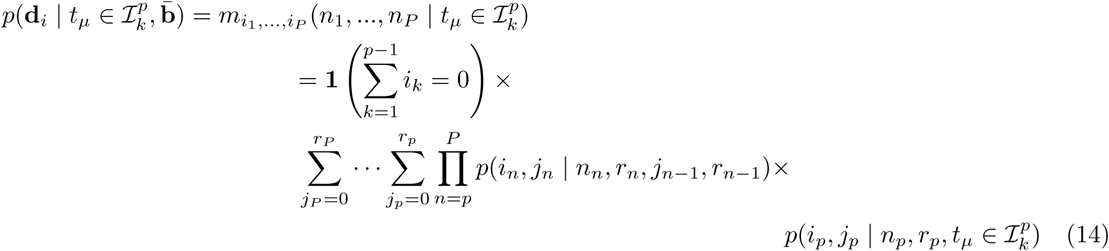

After running the MCMC inference, as described in [], we have samples from the posterior probability of 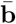. Given the *i*th MCMC sample 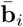, we calculate the probability of the mutation occurring in each of the intervals of 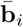 from (13), and then evaluate 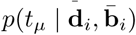 at a series of time points. We can then marginalise over 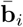’s to get the posterior density of *t*_*µ*_.

### Estimating the effective viral population size curve from NGS data

Given an estimate of the coalescent intervals of the genealogy underlying the JAFS, we estimate a model of effective viral population size over time, using the ‘skyline plot’ framework introduced in [22]. Skyline plot methods quantifies the relation between the genealogy of the sequences and the demographic history of the population, using coalescent theory concepts [23]. The coalescent framework describes the family relationships, or genealogy, between a set of sequences as a single realisation of a stochastic process in which pairs of lineages randomly coalesce, until all lineages can be traced backwards to a single common ancestor. The rate at which lineages coalesce along the genealogical timescale depends on the effective population size, enabling estimation of a demographic history of a set of sequences after estimating the genealogy.

We model the trajectory of viral population size with the Bayesian skyline plot [16], which allows uncertainty in the estimation of the coalescence times, and a piecewise constant model of population size change. Given the coalescence times of a set of sequences, the demographic history is described by *m* changes in population size through an ordered set of group sizes **A** = {*a*_1_, *…, a*_*m*_}, *a*_*i*_ *>* 0, 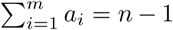 where *a*_*i*_ is the number of coalescent intervals in the *i*th group, and Θ = *θ*_1_, *…, θ*_*m*_, where *θ*_*i*_ is the effective population size within the *i*th interval.

For heterochronous data, as we consider in this paper, we introduce the terminology **u** = *u*_0_, *…, u*_*n*+*P*-2_, an ordered set of times, starting at the most recent sampled tips, and an indicator function *I*_*c*_(*i*) to indicate whether the *i*th event is a coalescent event (*I*_*c*_(*i*) = 1) or a sample event (*I*_*c*_(*i*) = 0). We use **w** to denote the times at which each grouped interval ends *w*_1_, *…, w*_*m*_, (a subset of **u**).

#### Simulating coalescent times under skyline model

The distribution of time to coalescence *t*_*c*_ for *n* lineages, given a variable effective population size of *θ*(*t*) is given by:

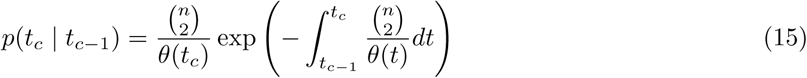

which depends on the average inverse population size over the interval, and the population size at the time of coalescence. Using the terminology detailed in the previous section, and the skyline model, and (15), we write the distribution of the *i*th interval Δ*u*_*i*_ = *|u*_*i*_ *-u*_*i-*1_*|* as:

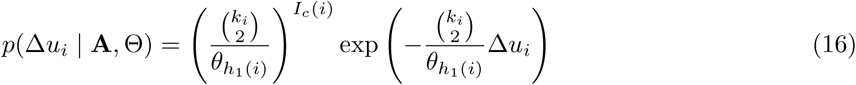

where *k*_*i*_ is the number of lineages in the *i*th interval, and *h*_1_(*i*) maps from indices in **u** to **w**:

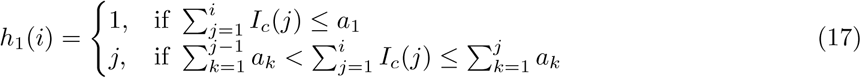

To simulate coalescent intervals under the skyline model, we use the following algorithm, based on inverse transform sampling. For ease of notation, we denote the coalescent times as **b** = *b*_1_,.., *b*_*N-*1_ (starting from most recent to oldest). **b** is a subset of **u**, 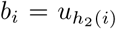, where *h*_2_(*i*) maps from indices in **u** to **b**: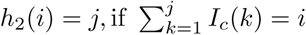, with *h*_0_(0) = 1.

With the *j*th set of tips sampled at time *t*_*j*_, *j* = *P*, …, 1, the cumulative density function of *b*_*i*_ is given by:

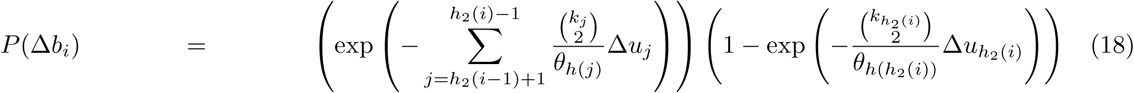

where Δ*b*_*i*_ is the *i*th coalescent interval,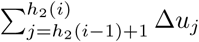.

After sampling *U ∼* Unif(0, 1), we find the inverse of (18) Δ*b*_*i*_ = *P* ^-1^(*U*). To sample from 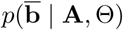, we average over *N*_*b*_ samples of Δ*b*_1_, *…,* Δ*b*_*N-*1_.

### MCMC implementation

We describe a method to sample the parameters of the Bayesian skyline plot model {Θ, **A**}, given the data *D*, the set of segregating site patterns. The posterior distribution is given by:

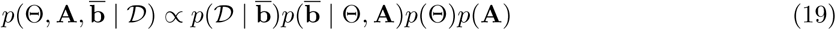

We use a two stage Metropolis Hastings proposal to sample from (19). A new value of {Θ^*∗*^, **A**^*∗*^} is proposed from *f* ({Θ^*∗*^, **A**^*∗*^} *|* {Θ, **A**}), and then a corresponding 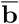 is sampled from 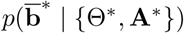. The proposal 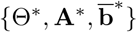 is accepted with probability

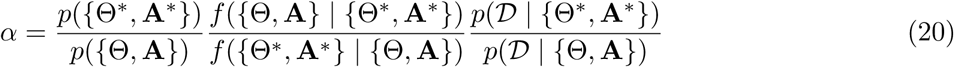

## Supporting Information

### Model of viral dynamics

We describe a model of viral dynamics [3, 24, 25]. In this model, influenza A virus infection is limited by the availability of susceptible target (epithelial) cells rather than the effects of the immune response, and virus production is delayed. This is described by the following differential equations:

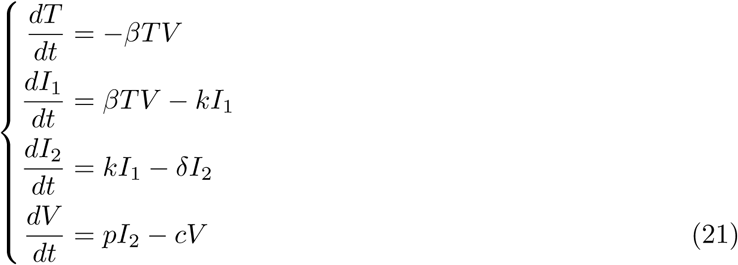

*T* is the number of uninfected target cells. The delay in the production of free virus is modelled by defining two separate populations of infected epithelial cells; one population *I*_1_ is infected but not yet producing virus; the second population *I*_2_ is actively producing virus. The average transition time between *I*_1_ and *I*_2_ is given by 1*/k*. *V* represents the viral population. Susceptible cells become infected by virus at rate *βTV*, where *β* is the rate constant characterising infection. Virally infected cells, *I*_2_, by shedding virus, increase the viral population at an average rate of *p* per cell and die at a rate of *δ* per cell. Free virus is cleared at a rate of *c* per day.

The units in which *V* is measured will affect the scaling of the model parameters. For instance, *V* represents the infectious viral titer expressed in TCID_50_*/*ml of nasal wash in [3], whereas in [4], *V* is the number of RNA copies/ml. In this work, we choose *V* to represent the number of virions in the population, so that each member of the population represents a viral genome. Following [17], in which the rate of coalescence was derived for a deterministic SIR compartmental model, the rate of coalescence in a population of *n* under the Baccam model is given by:

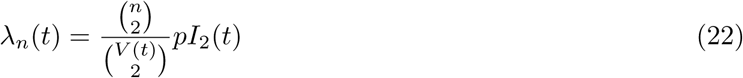

By creating a piecewise constant approximation of *V* (*t*), and *I*_2_(*t*), with Δ*t* = 1*e*^−4^ days, we can simulate coalescent intervals as in Section () (note that we do not necessarily end with 1 lineage at *t* = 0).

For comparison with the skyline approximation to the effective population size trajectory that we note that:

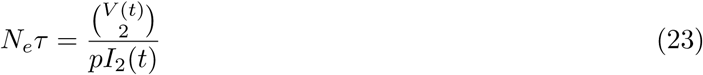

where *τ* is the (unknown) viral generation time in days.

Average values of the parameters are estimated in [4], which are displayed in Table 1.

## Acknowledgments

The authors wish to thank Pablo Murcia and Joseph Hughes at the Centre for Virus Research, University of Glasgow, for permission to use their equine influenza dataset, and for helpful comments and advice. The work was funded by the Wellcome Trust [WR/094527].

